# Light-induced difference FTIR spectroscopy of primate blue-sensitive visual pigment at 163 K

**DOI:** 10.1101/2020.05.29.123224

**Authors:** Shunpei Hanai, Kota Katayama, Hiroo Imai, Hideki Kandori

## Abstract

Structural studies of color visual pigments lag far behind those of rhodopsin for scotopic vision. Using difference FTIR spectroscopy at 77 K, we report the first structural data of three primate color visual pigments, monkey red (MR), green (MG), and blue (MB), where the batho-intermediate (Batho) exhibits photoequilibrium with the unphotolyzed state. This photochromic property is highly advantageous for limited samples since the signal-to-noise ratio is improved, but may not be applicable to late intermediates, because of large structural changes to proteins. Here we report the photochromic property of MB at 163 K, where the BL intermediate, formed by the relaxation of Batho, is in photoequilibrium with the initial MB state. A comparison of the difference FTIR spectra at 77 and 163 K provided information on what happens in the process of transition from Batho to BL in MB. The coupled C_11_=C_12_ HOOP vibration in the planer structure in MB is decoupled by distortion in Batho after retinal photoisomerization, but returns to the coupled C_11_=C_12_ HOOP vibration in the *all-trans* chromophore in BL. This suggests that BL harbors a planer all-*trans* configuration of retinal. The Batho formation accompanies helical structural perturbation, which is relaxed in BL. The H-D unexchangeable X-H stretch weakens the hydrogen bond in Batho, but strengthens it in BL. Protein-bound water molecules that form an extended water cluster near the retinal chromophore change hydrogen bonds differently for Batho and BL, being stronger in the latter than in the initial state. In addition to structural dynamics, the present FTIR spectra at 163 K show no signals of protonated carboxylic acids as well as 77 K, suggesting that E181 is deprotonated in MB, Batho and BL.

## INTRODUCTION

Humans have two kinds of vision: twilight vision mediated by rhodopsin in rod photoreceptor cells and color vision achieved by multiple color pigments in cone photoreceptor cells [1]. Humans possess three color pigments: red-, green-, and blue-sensitive proteins maximally absorbing at 560, 530, and 425 nm, respectively [2]. Rhodopsin and color-pigments both contain a common chromophore molecule, 11-*cis* retinal, whereas different chromophore-protein interactions allow preferential absorption of different colors [3, 4]. Purified proteins are a prerequisite to understand the mechanism of color tuning and activation process by light. Studying rhodopsin is highly advantageous because large amounts of protein can be obtained from vertebrate and invertebrate native cells. Consequently, X-ray structures were determined not only for the unphotolyzed state of bovine [5], squid [6], and jumping spider [7] rhodopsins, but also for opsin [8], photobleaching intermediates [9–12], active state [13,14] and the active-state complexed with the C-terminus peptide of the α subunit of G-protein [14–16] or engineered mini-Go [17], and arrestin [18]. These structures have provided insight into the mechanism of the chromophore-protein interaction and activation.

On the other hand, structural studies of color pigments lag far behind those of rhodopsin. No color visual pigments have been crystallized. The structural analysis of green and red pigments was only reported by resonance Raman spectroscopy, in which the observed vibrational bands were very similar between human green and red, indicating similar chromophore-protein interactions [19]. It should be noted that resonance Raman spectroscopy only provides vibrational signals from the chromophore, but not from the protein. The involvement of water dipoles was discussed [19], but no experimental confirmation was possible for protein-bound water molecules. In contrast, IR spectroscopy is able to provide vibrational signals not only from the chromophore, but also from protein and water molecules. We initiated light-induced difference FTIR spectroscopy of monkey red- (MR), monkey green- (MG), and monkey blue (MB)-sensitive color visual pigments, which were expressed in HEK cells.

Preparation of a hydrated film sample needs 0.1-0.15 mg purified protein, which corresponds to about 100 dishes 10 cm in diameter. Consequently, we have reported structural information for MR and MG since 2010 [20–26]. Although a lower expression level was reported for blue-sensitive pigments, we successfully obtained light-induced difference FTIR spectroscopy of MB from marmoset [27].

Except for a chloride-binding study by ATR-FTIR spectroscopy at room temperature [25, 26], all reported measurements were performed at 77 K, where the photoproduct, batho-intermediate (Batho), can be reverted to the initial state by light. Consequently, photoconversions from the initial state to Batho, and Batho to the initial state were repeated. This photochromic property has been highly advantageous to improve the signal-to-noise ratio of FTIR spectra. It is believed that this photochromic property is unique to 77 K, and is not applicable to late intermediates. In fact, in the case of bovine rhodopsin, illumination of the lumi-intermediate (Lumi), which is stable at 200 K, is not reverted to the initial state [28], presumably because of large structural changes to the protein. Thus, although difference FTIR spectroscopy is a powerful method to study color visual pigments, such studies have been limited to the primary intermediate, Batho.

Here we report that MB performs the above described photochromic property not only at 77 K, but also at 163 K, where the BL intermediate is formed by the relaxation of Batho. By using this advantage, we accumulated difference FTIR spectra for BL, and the spectra obtained were compared to those of Batho. We found a highly distorted C_11_=C_12_ bond in Batho, but becoming planer in BL. A helical structural perturbation in Batho is also relaxed in BL. In contrast, protein-bound water molecules that form an extended water cluster near the retinal chromophore change hydrogen bonds differently for Batho and BL, stronger in the latter than in the initial state. The molecular mechanism of the light-induced activation process in MB is discussed.

## MATERIALS AND METHODS

### Sample Preparation

MB cDNA was tagged with the Rho1D4 epitope sequence and introduced into the pFastBac HT expression vector. This construct was expressed in the Sf9 cell line and regenerated with 11-*cis*-retinal [25, 27]. The regenerated sample was solubilized with a buffer containing 2% (w/v) *n*-dodecyl-β-D-maltoside (DDM) (final concentration was 1% (w/v)), 50 mM HEPES, 140 mM NaCl, and 3 mM MgCl_2_ (pH 7.0) and purified by adsorption on an antibody-conjugated column and eluted with a buffer containing 0.10 mg/mL 1D4 peptide, 0.02% DDM, 50 mM HEPES, 140 mM NaCl, and 3 mM MgCl_2_ (pH 7.0).

### Low-Temperature UV-visible and FTIR Spectroscopy

Low-temperature UV-visible spectroscopy was applied to the films hydrated with H_2_O at 77 K as described previously [29, 30]. Low-temperature FTIR spectroscopy was applied to the films hydrated with H_2_O, D_2_O, or D2^18^O at 163 K, as described previously [20, 21, 27]. For the formation of the Batho intermediate of MB, samples were irradiated with 400 nm light (by using an interference filter) for 5 min. For the reversion from the Batho intermediate to the dark state, samples were irradiated with >480 nm light for 5 min. For each measurement, 128 interferograms were accumulated, and 40 recordings were averaged. FTIR spectra were recorded with 2 cm^−1^ resolution.

## RESULTS

### Photochromism between MB and the BL Intermediate at 163 K

The blue curve in Figure 1a shows the absorption spectrum of MB at 77 K, and illumination of MB with 400-nm light yielded a spectral red-shift (red curve), indicating the formation of Batho. Subsequent illumination at >480 nm recovered the original absorption spectrum, which is evident from the mirror-imaged difference spectra (Figure 1b). The photochromic property for primary intermediates is well-known for rhodopsins, not only for animal rhodopsins such as MB, but also for microbial rhodopsins [3]. The structure of the retinal binding pocket changes little at cryogenic temperatures such as 77 K [9, 11], so that retinal photoisomerization is reversible, between the 11-*cis* and *all-trans* forms in animal rhodopsins, and between the *all-trans* and 13-*cis* forms in microbial rhodopsins.

**Figure 1:**
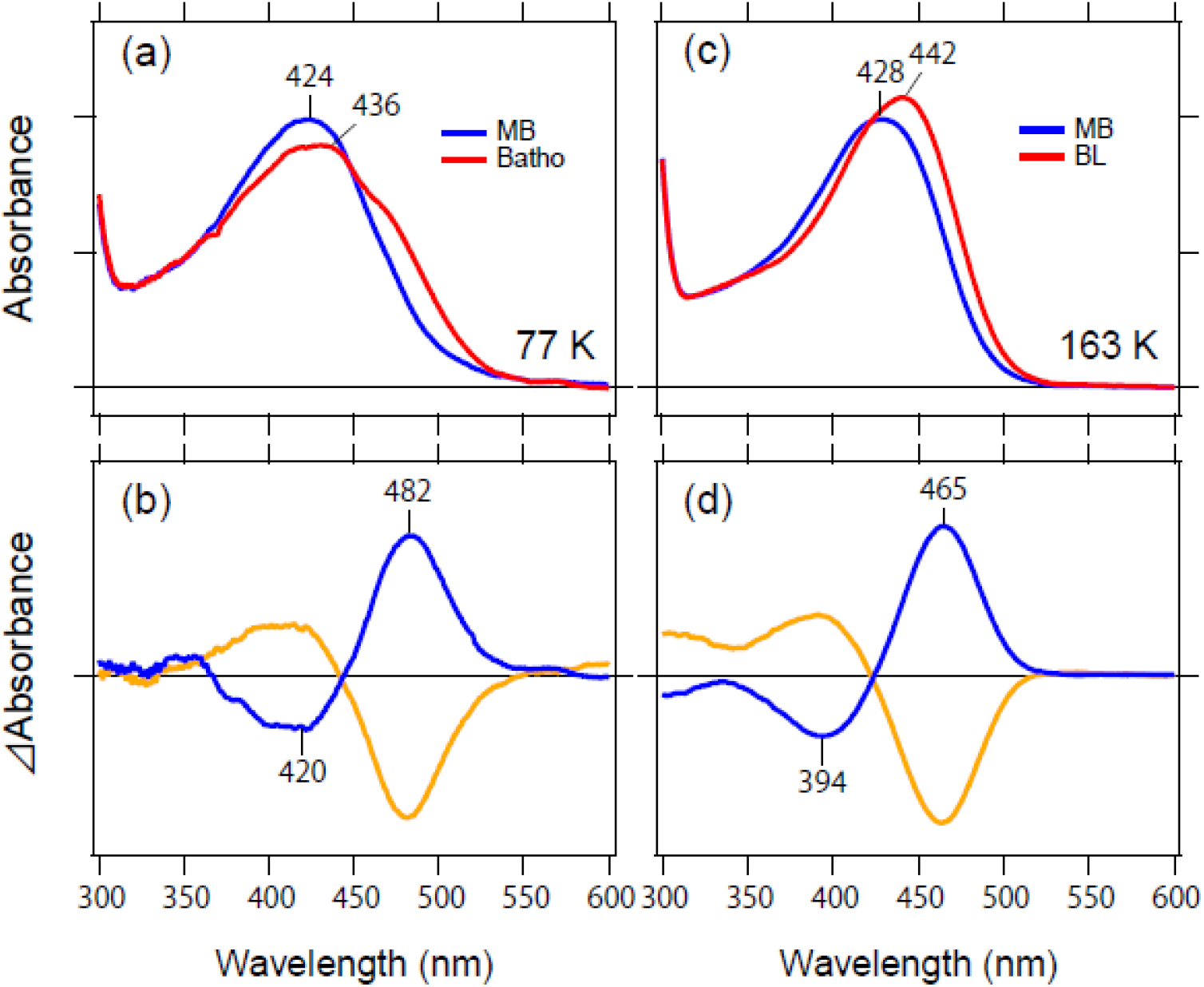
(a) Absorption spectra of monkey blue (MB) (blue curve) at 77 K, and photoequilibrium under illumination at 400 nm (red curve). Subsequent illumination at >480 nm results in the initial spectrum (blue curve). (b) Light-induced difference spectra by 400-nm illumination (blue curve) and by >480-nm illumination (orange curve). (c) Absorption spectra of MB (blue curve) at 163 K, and photoequilibrium under illumination at 400 nm (red curve). Subsequent illumination at >480 nm results in the initial spectrum (blue curve). (b) Light-induced difference spectra by 400-nm illumination (blue curve) and by >480-nm illumination (orange curve).

This photochromic property is not applicable to late intermediates, where protein structural changes take place. In the case of bovine rhodopsin, the subsequent intermediate after Batho is Lumi, meta-rhodopsin I (Meta I), and meta-rhodopsin II (Meta II), which can be trapped at 200 K, 240 K, and 270 K, respectively. None of these exhibit the photochromic property, unlike Batho, presumably because of structural changes in each intermediate [28]. A previous study of chicken blue (CB)-sensitive pigment revealed the presence of the BL intermediate between Batho and Lumi [31]. The transition temperatures from Batho to BL, and from BL to Lumi were 113 K and 163 K, respectively, but their photochromic properties were not studied previously. In our study, we found that the transition temperature from Batho was located at 120-130 K, which is stable up to 170 K. We postulate the intermediate as BL of MB from the analogy to CB [31]. The blue curve in Figure 1c shows the absorption spectrum of MB at 163 K, and illumination of MB by 400-nm light yielded a spectral red-shift (red curve). Subsequent illumination at >480 nm recovered the original absorption spectrum, which is evident from the mirror-imaged difference spectra (Figure 1d). The positive peak at 465 nm was blue-shifted by 20 nm from that at 77 K (Figure 1b). These results indicate the photochromic property of the BL intermediate of MB at 163 K.

Difference FTIR spectra in Figure 2 correspond to those in UV-visible (Figure 1d) at 163 K. Prominent peaks were observed for the frequency region of the C=C stretch (1600-1550 cm^−1^), the C-C stretch (1250-1150 cm^−1^), and hydrogen out-of-plane (HOOP) vibrations (1000-800 cm^−1^) of the retinal chromophore. The photochromic property of MB at 163 K is highly advantageous for structural analysis when using difference FTIR spectroscopy, as the signal-to-noise ratio can be improved by repeated measurements. Consequently, we accumulated the spectra at 163 K as well as at 77 K, both of which are compared next.

**Figure 2:**
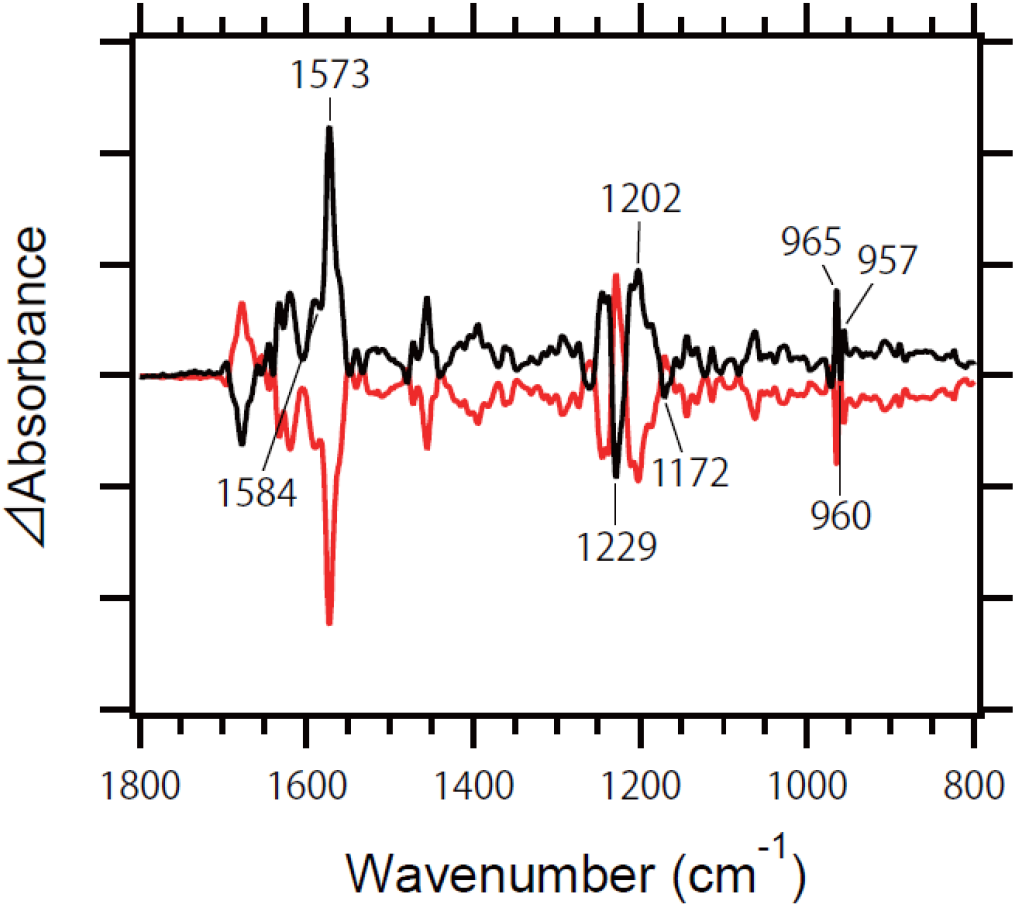
Light-induced difference FTIR spectra of monkey blue (MB) at 163 K. The spectra were obtained by 400-nm illumination (black curve) and by >480-nm illumination (orange curve), which correspond to the BL-minus-MB and MB-minus-BL spectra, respectively. One division of the y-axis corresponds to 0.0015 absorbance units.

### Comparison of Light-Induced Difference FTIR Spectra of MB at 163 L and 77 K. (1) Chromophore Vibrations

Figure 3 compares the light-induced difference FTIR spectra (light-minus-dark) of MB at 163 K and 77 K, which were measured in H_2_O (black curve) and D_2_O (red curve). At 77 K, a clear peak pair was observed at 1576 (-)/1560 (+) cm^−1^, which corresponds to the ethylenic C=C stretching vibration of MB and Batho. The spectral down-shift in this vibrational mode of retinal chromophore at 1580-1500 cm^−1^ corresponds to the red-shift in the visible region. At 163 K, a strong positive peak was observed at 1573 cm^−1^, whereas the corresponding negative peak was unclear. We infer that the peak pair at 1584 (-)/1573 (+) cm^−1^ corresponds to the ethylenic C=C stretching vibration of MB and the intermediate at 163 K (BL), where a broad positive band is superimposed. The positive peaks at 1573 and 1560 cm^−1^ at 163 K and 77 K, respectively, are consistent with the blue-shifted UV-visible absorption of BL. The negative peak from the ethylenic C=C stretch of MB should be identical, but the apparent high frequency shift of the negative peak at 163 K (1584 cm^−1^) from that at 77 K (1575 cm^−1^) presumably originates from the high frequency shift of the positive peak.

**Figure 3:**
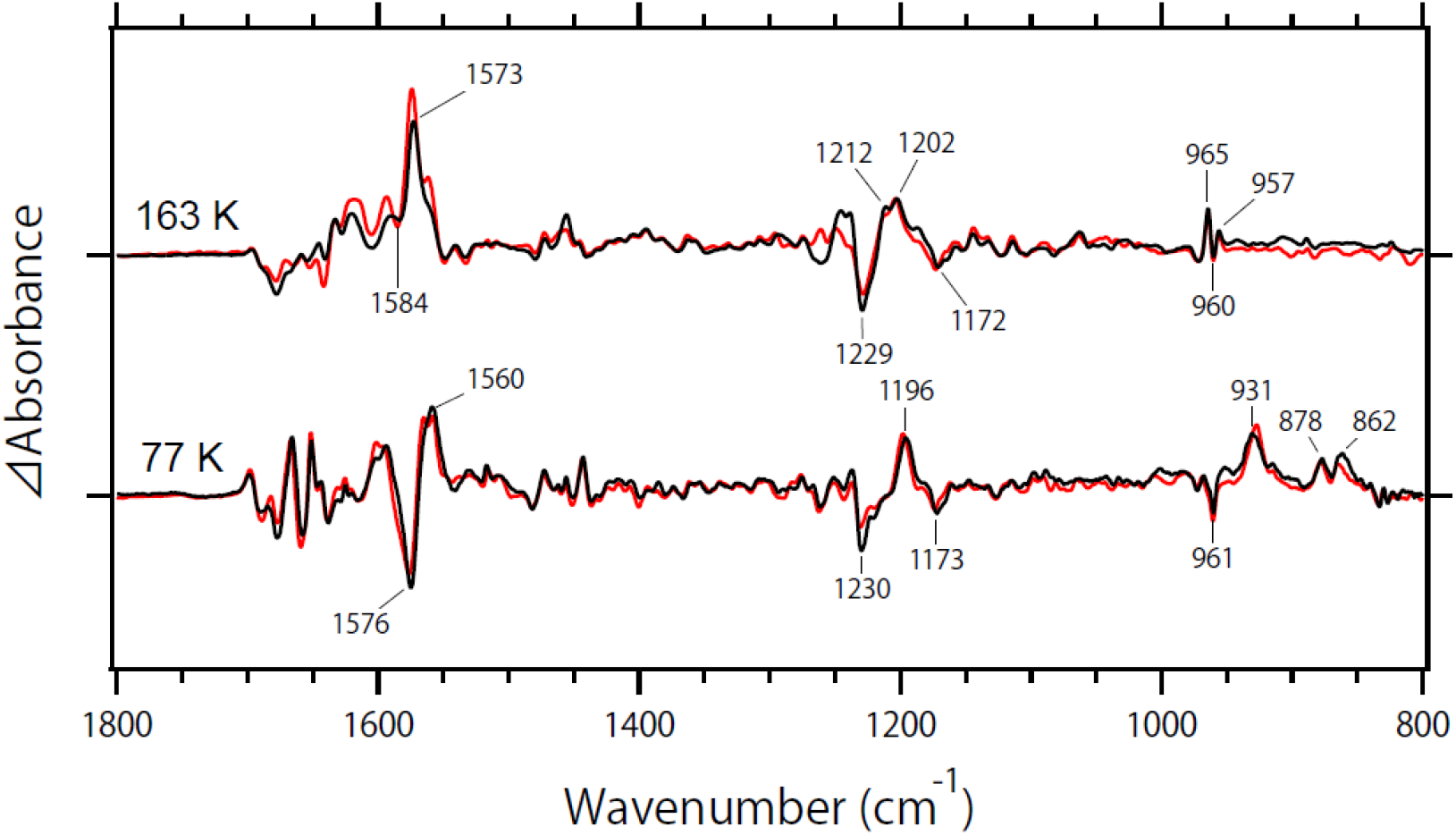
Light-minus-dark difference FTIR spectra of MB at 163 K and 77 K in the 1800-800 cm^−1^ region. Spectra were measured in H_2_O (black curve) and D_2_O (red curve). Negative bands originate from MB, while positive bands originate from the BL-intermediate (BL) and batho-intermediate (Batho) at 163 K and 77 K, respectively. One division of the y-axis corresponds to 0.004 absorbance units.

The frequency region at 1250-1150 cm^−1^ exhibits the C-C stretching vibrations of retinal chromophore (Figure 3). While the negative peaks at 1229/1230 and 1172/1173 cm^−1^ for MB are common at 163 and 77 K, the positive peak is different between 163 K and 77 K, suggesting that the retinal conformation differs between Batho and BL. At 77 K, a single positive peak was observed at 1196 cm^−1^. In contrast, two peaks were observed at 1212 and 1202 cm^−1^ at 163 K. These results probably accompany relaxation of the retinal chromophore from Batho to BL.

The frequency region at 1000-800 cm^−1^ exhibits the HOOP wagging mode of retinal chromophore and reflects a structurally perturbed and/or distorted chromophore (Figure 3; highlighted in Figure 4). At 77 K, a sharp negative peak appeared at 961 cm^−1^, and a similar band was observed at 969 cm^−1^ in bovine rhodopsin, which was assigned as the C_11_=C_12_ HOOP modes [32]. Upon formation of Batho, multiple intense peaks were observed at 931, 878, and 862 cm^−1^, and enhanced HOOP vibrations indicate a highly distorted chromophore structure after 11-*cis* to *all-trans* retinal isomerization in the restricted protein environment. The C_11_H and C_12_H wagging modes for Batho of bovine rhodopsin were reported at 921 cm^−1^ and 858 cm^−1^, respectively [32–34]. While these band assignments were performed by resonance Raman spectroscopy, similar band assignment was reported for chicken red (iodopsin) by using difference FTIR spectroscopy at 77 K [35]. From this analogy, the peaks at 931 and 862 cm^−1^ are ascribable for the C_11_H and C_12_H wagging modes of Batho of MB. Slight spectral shifts were observed for these bands in D_2_O, suggesting that other vibrations influenced by H-D exchange may be involved.

**Figure 4:**
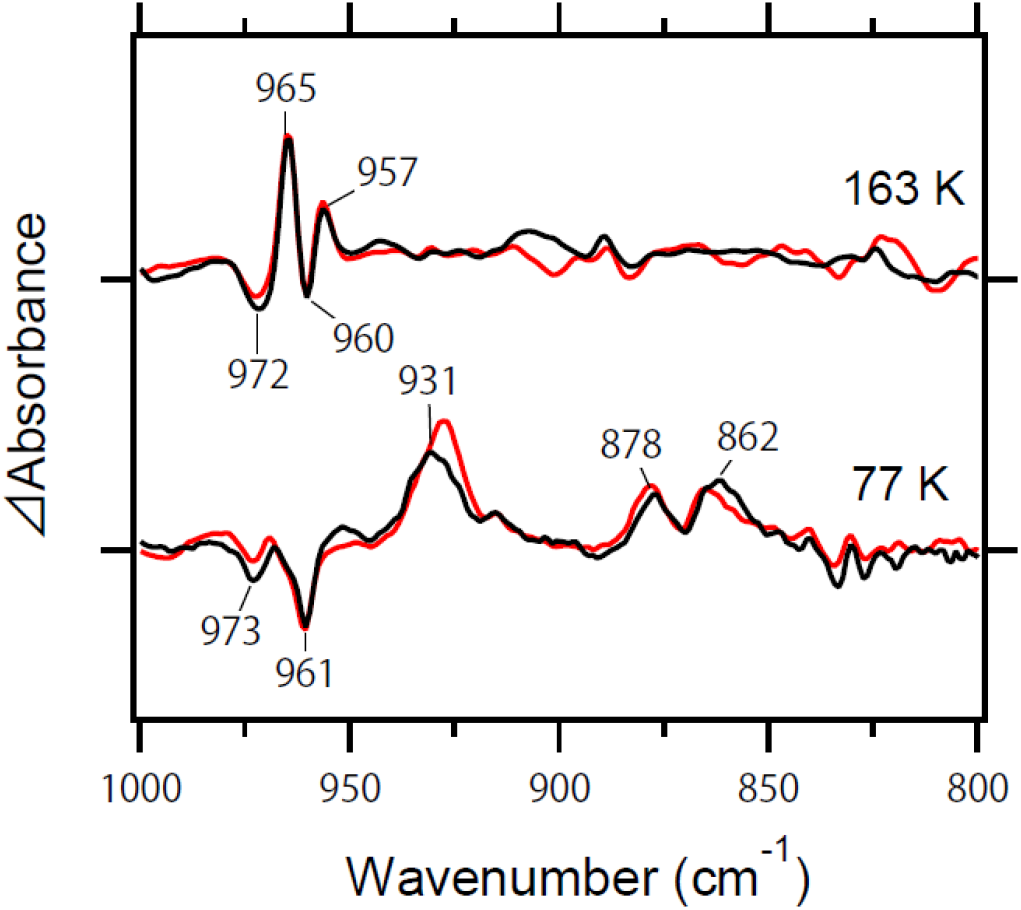
Expanded spectra from Figure 3 in the 1000-800 cm^−1^ region. One division of the y-axis corresponds to 0.004 absorbance units. Spectra were measured in H_2_O (black curve) and D_2_O (red curve).

A very different spectral feature was observed at 163 K, where intense peaks at 930-850 cm^−1^ almost disappeared. Instead, sharp positive peaks appeared at 965 and 957 cm^−1^, which correspond to the negative peaks at 972 and 960 cm^−1^. The negative 960-cm^−1^ band probably originates from the coupled C_11_=C_12_ HOOP modes, while the origin of the negative 972-cm^−1^ band is unclear at present. As there is no effect by deuterium, the C_14_H and C_15_H wagging modes are unlikely. Therefore, HOOP vibrations such as C_7_H, C_8_H and C_10_H wagging modes are candidates.

The band shape of the coupled C_11_=C_12_ HOOP mode is sharper in MB (at 960 cm^−1^) than in MG and MR [27], implying that C_11_=C_12_ retains its geometric planar structure compared to other pigments in their dark states. Upon retinal photoisomerization, the strongly coupled double bond at C_11_ and C_12_ of the retinal polyene chain are decoupled, leading to C_11_H and C_12_H wagging modes at 928 and 863 cm^−1^, respectively, in Batho. The transition from Batho to BL regains its coupled C_11_=C_12_ HOOP mode, whose frequency (965 or 957 cm^−1^) is similar to that in MB (960 cm^−1^). This suggests that BL contains the planer C_11_=C_12_ group. Interestingly, BL contains the all-*trans* chromophore as well as Batho, but the local distortion of chromophore is similar to MB that contains the 11-*cis* chromophore.

### Comparison of Light-Induced Difference FTIR Spectra of MB at 163 L and 77 K. (2) Protein Vibrations

Figure 5a shows the difference FTIR spectra at 1780-1620 cm^−1^ that mainly monitors vibrations of proteins such as the C=O stretch of protonated carboxylic acids and the peptide backbone (amide-I vibration) at 1780-1700 cm^−1^ and 1700-1620 cm^−1^, respectively. No spectral changes at >1700 cm^−1^ at 77 K indicates no changes to hydrogen bonding of protonated carboxylic acids upon retinal photoisomerization. This is also the case at 163 K, indicating that protonated carboxylic acids are not involved in structural changes upon formation of BL. Figure 6 shows four carboxylates, D83, E113, E122 and E181, in bovine rhodopsin, and the corresponding amino acids in primate color pigments. It is well established that E113 is deprotonated as the counterion of the protonated Schiff base, while D83 and E122 are protonated in bovine rhodopsin, whose signals are clearly observable at 1780-1700 cm^−1^ [36–38]. In contrast, there is a continuing debate whether E181 is protonated or not [39–44]. MB contains E113 and E181, where E113 is deprotonated as the Schiff base counterion. No signal at >1700 cm^−1^, not only at 77 K, but also at 163 K, suggests that E181 is deprotonated.

**Figure 5:**
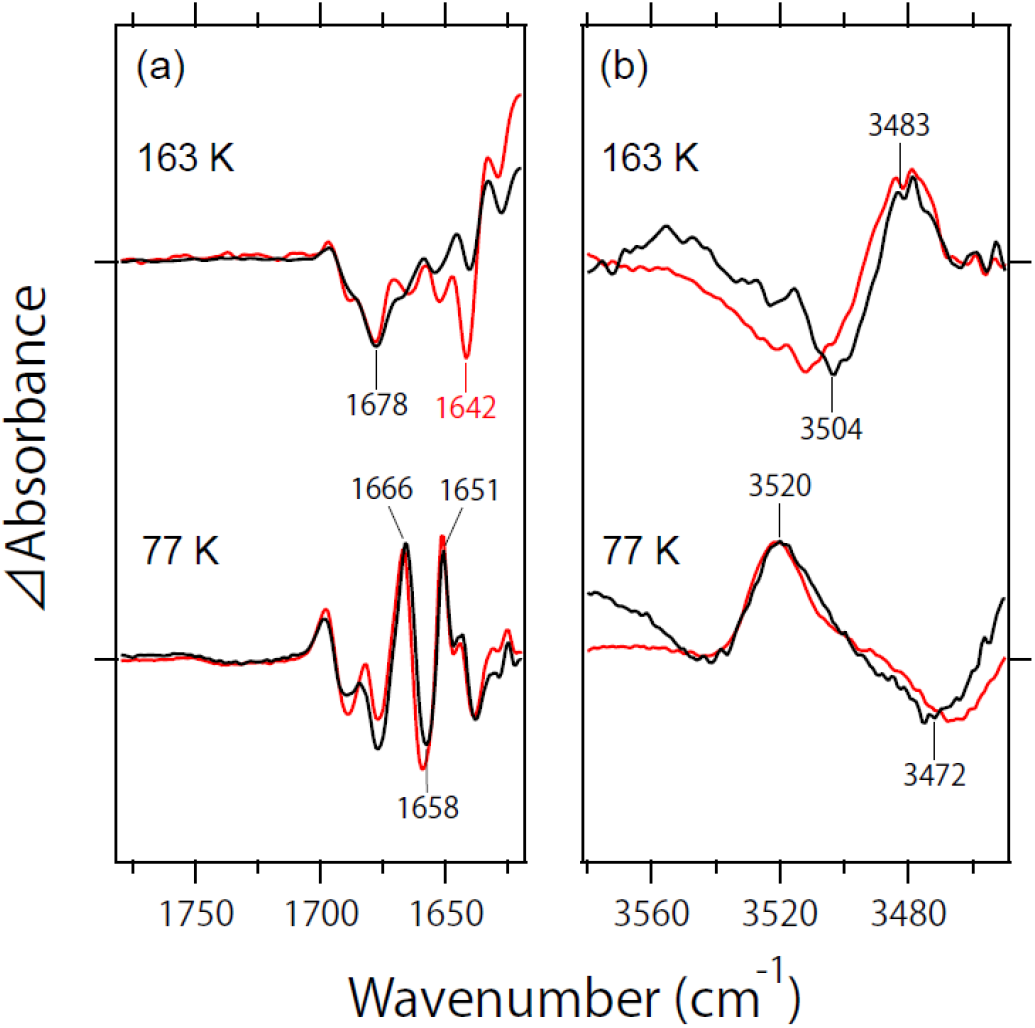
(a) Expanded spectra from Figure 3 in the 1780-1620 cm^−1^ region. One division of the y-axis corresponds to 0.006 absorbance units. Spectra were measured in H_2_O (black curve) and D_2_O (red curve). (b) Light-minus-dark difference FTIR spectra of MB at 163 K and 77 K in the 3580-3450 cm^−1^ region.

**Figure 6:**
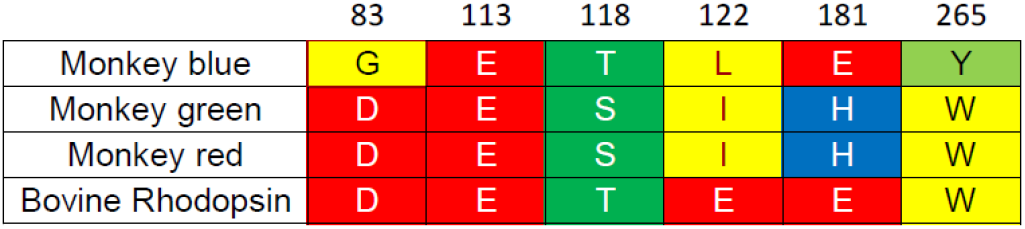
Comparison of key residues in MB, MG, MR and bovine rhodopsin.

Amide-I vibrations appear at 1700-1620 cm^−1^ as well as vibrations of side chains. It should be noted that the C=N stretch of the retinal chromophore is also involved in this frequency region, whose frequency differs between H_2_O (C=NH) and D_2_O (C=ND). At 77 K, two spectra in H_2_O and D_2_O are superimposable with each other, suggesting no frequency changes of the C=N vibration between MB and Batho. This is also the case for bovine rhodopsin and Batho, where an alteration in structure occurs only the middle of the chromophore whereas the hydrogen bond of the Schiff base is unchanged. At 163 K, we observed a negative peak at 1642 cm^−1^ and a broad positive band centered at 1618 cm^−1^. The 1642-cm^−1^ band may originate from the C=ND stretch of the chromophore, although the corresponding C=NH stretch is unclear. However, if it is, then the hydrogen bond of the Schiff base is preserved in Batho, but altered in BL.

As we reported previously, MB shows stronger peaks in the amide-I frequency of the α-helix than MG and MR at 77 K [27], suggesting the largest helical structural changes upon retinal photoisomerization in MB. Figure 5a shows peaks of amide-I at 1676 (-)/1666 (+)/1658 (-)/1651 (+) cm^−1^ upon formation of Batho at 77 K, among which the negative peak at 1658 cm^−1^ corresponds to the amide-I frequency of theα-helix. Interestingly, large structural changes in α-helices disappear upon formation of BL. Relaxation of helical structural changes in the transition from Batho to BL takes place together with relaxation of the distorted C_11_=C_12_ group. On the other hand, the negative band at 1676 cm^−1^ seems commonly present at 77 and 163 K.

Under D_2_O hydration, the vibrational band at 4000-2700 cm^−1^ (X-H stretching vibration) originates from H-D unexchangeable O-H and N-H stretching vibrations of amino acids, which are involved in the hydrophobic environment. One of the characteristic bands at 3550-3450 cm^−1^ is the O-H stretch of Thr or Ser, which was first observed for bovine rhodopsin [44]. In bovine rhodopsin, the O-H group of T118, whose stretching frequency is located at 3463 cm^−1^ [45], is only deuterated at high temperatures [46], which was used to investigate protein fluctuation as the origin of dark noise [47–50]. A previous study suggested that the bands at 3520 (+)/3472 (-) cm^−1^ are candidates of the O-H stretch of T118 in MB [27], where a high frequency shift indicates a weaker hydrogen bond upon formation of Batho. Interestingly, the frequency shift (3504 (-)/3483 (+) cm^−1^) was opposite at 163 K (Figure 5b), where the hydrogen bond is stronger in BL than in MB. Thus, based on the postulation of the O-H stretch of T118, the following interpretation is possible. The hydrogen bond is weakened upon Batho formation, together with the distortion of the C_11_=C_12_ group in the all-*trans* chromophore and helical structural perturbation. Then, the O-H group of T118 finds the hydrogen-bonding acceptor in BL, whose hydrogen bond is stronger than in MB, along with the relaxation of the C_11_=C_12_ distortion and helical structural perturbation.

### Comparison of Light-Induced Difference FTIR Spectra of MB at 163 L and 77 K. (3) Water Vibrations

We previously reported unique water vibrations for MB. As is seen in Figure 7, a very broad and high intensity positive peak appears in the 2600-2500 cm^−1^ region at 77 K [27], whose frequency corresponds to deuterated water ice in a tetrahedral hydrogen bonding network. As the broad and intense band was influenced by the mutation of Y265 and E113, we proposed the presence of an extended water cluster between Y265 and E113. Y265 is specific to blue-sensitive pigments [27] while other pigments, including rhodopsin, contain W265 that plays an important functional role. From the similar frequencies at 2567 and 2494 cm^−1^, an extended water cluster between Y265 and E113 seems to be present in the unphotolyzed state of MB.

**Figure 7:**
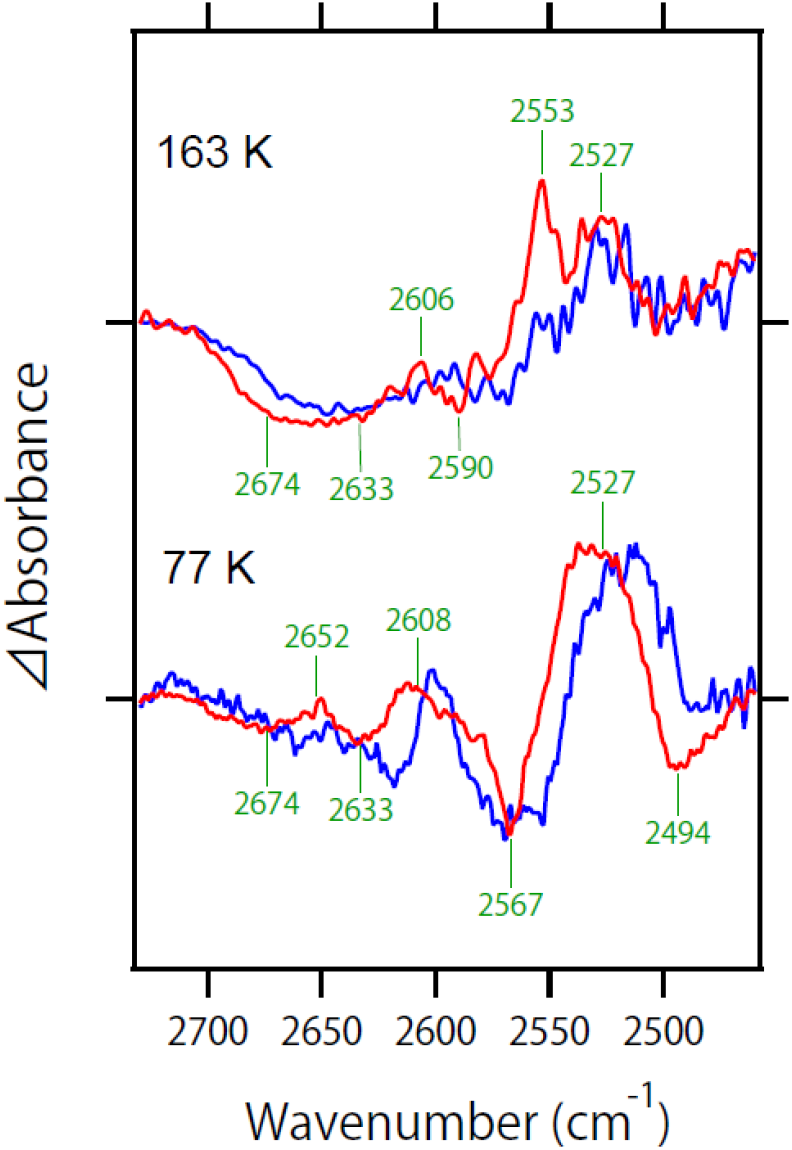
Light-minus-dark difference FTIR spectra of MB at 163 K and 77 K in the 2730-2450 cm^−1^ region. Spectra were measured in D_2_O (red curve) and D2^18^O (blue curve).

At 163 K, observed water signals are similar to those at 77 K, with some variations (Figure 7). At 77 K, two bands at 2674 (-)/2652 (+) cm^−1^ and 2633 (-)/2608 (+) cm^−1^ look peak pair with each other. Two negative water O-D stretches are preserved at 2674 and 2633 cm^−1^ in the 163 K spectra, but disappearance of the positive band at 2652 cm^−1^ resulted in the broad negative water band at 2700-2600 cm^−1^. Corresponding positive bands of the water O-D stretch appear at 2600-2500 cm^−1^. Thus, it is likely that the transition from Batho to BL accompanies rearrangement of water molecules in the cluster, where the hydrogen bond is stronger than in MB.

## DISCUSSION

Light-induced difference FTIR spectroscopy is a powerful method to study the structural dynamics of photoreceptive proteins such as rhodopsins. However, in the case of animal rhodopsins, the photobleaching property made this difficult since the measurements from a single sample cannot be repeated. The only exception is the primary intermediate, Batho, which is in photoequilibrium with the initial state [45]. Thus, the difference FTIR spectra at 77 K, which have a high signal-to-noise ratio by repeating the measurements, provided important structural information about the initial and Batho states in bovine Rh, monkey Rh, MG, MG, and MB [20, 27, 45]. In the present study, we report a similar photochromic property for the BL intermediate of MB at 163 K, which allowed us to repeat the measurements, leading to accurate spectral acquisition. The reason for the successful spectral acquisition of MB is unclear at present. In the case of bovine rhodopsin, Batho is photoconverted to the initial state, but Lumi is not [28].

Existence of the blue-shifted intermediate (BSI) was reported between Batho and Lumi for bovine rhodopsin [49, 50], whereas BSI cannot be stabilized at low temperature. The BL intermediate in MB may correspond to BSI, which can be stabilized at 163 K. This could be the reason.

The obtained difference spectra for BL at 163 K were compared with those for Batho at 77 K, which provided useful information on how light energy stored in the structure of Batho is relaxed and transferred in BL. Analysis of the HOOP vibration revealed that the C_11_=C_12_ *cis* form of the retinal polyene chain is locally planer in the initial state, but highly distorted in the *trans* form of Batho, while the transition from Batho to BL yields a planer C_11_=C_12_ group in the all-*trans* form. Upon the formation of Batho, helical structural changes take place, which recovers in BL, suggesting that retinal isomerization causes local helical perturbation, but relaxes in BL. The environment around T118 of MB seems to be similar to rhodopsin because of the almost same O-H stretching frequency (3463 cm^−1^) [46, 47], suggesting that the O-H group forms a hydrogen bond with the peptide carbonyl of G114 in MB. The hydrogen bond is weakened in Batho, but is further strengthened in BL. T118 may find a different hydrogen-bonding acceptor. We proposed an extended water cluster between Y265 and E113 in MB [27], which is locally perturbed in Batho, and further rearranged in BL, where hydrogen bonds are much stronger than in the initial state.

Vibrational analysis of the amide-I band showed that helical structural changes take place in Batho, but such changes diminish in BL. This does not necessarily mean that MB and BL have the same structure. Amide-I vibrational frequency is highly sensitive to the secondary structure of the peptide backbone, so helical structural changes such as kink motion and deformation can be monitored with high sensitivity. In contrast, the amide-I vibration is insensitive to helix motion without altering the secondary structure, such as translational motion. Therefore, it is possible that helical structural changes in Batho are transferred to a different arrangement of helices.

E181 in bovine rhodopsin is an important residue in the animal rhodopsin family. It is the Schiff base counterion in invertebrate rhodopsins, and it was proposed that ancestral vertebrate rhodopsin acquired a new counterion at position 113 during molecular evolution [53]. Interestingly, vertebrate rhodopsins preserve E181, whose protonation states have long been debated [39–44]. The 1780-1700 cm^−1^ region in difference FTIR spectroscopy is good at monitoring protonated carboxylic acids, and the spectral analysis of bovine rhodopsin suggested no signal owing to E181 [36–38, 40]. This observation can be interpreted by (i) E181 is deprotonated, or (ii) E181 is protonated, but without changes in the hydrogen bond between the initial and intermediate states. In spite of many experimental and theoretical efforts [36–44], the protonation state of E181 has not yet been concluded in bovine rhodopsin. Blue-sensitive pigments such as MB also contain E181, and a previous study [27] as well as the present study showed no signal at >1700 cm^−1^ both at 77 K and 163 K, respectively. This observation for MB can be interpreted by (i) E181 is deprotonated, or (ii) E181 is protonated, but without changes in the hydrogen bond between MB, Batho, and BL. In the analysis of bovine rhodopsin, signals from D83 and E122 appear in the 1800-1700 cm^−1^ region, which complicates the analysis of E181. In contrast, MB contains neither D83 nor E122 (Figure 6), and it is much easier to analyze the data in the 1780-1700 cm^−1^ region [36–38, 40]. Further studies on Lumi and Meta-I of MB will unequivocally reveal the protonation state of E181.

Light energy is stored in protein by retinal photoisomerization, where chromophore distortion plays a key role. Then, energy transfer from the distorted chromophore is important to activate visual pigments, one type of G-protein coupled receptors. A recent study reported that one microbial rhodopsin exhibits retinal photoisomerization, but the primary K intermediate returns to the initial state without undergoing subsequent reactions [30]. Therefore, proper energy transfer is needed, and the present study revealed the participation of a protein moiety and protein-bound water molecules.

## CONCLUSION

We found the photochromic property of MB at 163 K, and not only at 77 K, which enabled a highly accurate spectral acquisition for BL and Batho. A comparison of difference FTIR spectra at 77 K and 163 K leads to the events from Batho to BL. The coupled C_11_=C_12_ HOOP vibration in the planer structure in MB is decoupled by a distortion in Batho after retinal photoisomerization, but returns to the coupled C_11_=C_12_ HOOP vibration in the all-*trans* chromophore in BL. Batho formation accompanies helical structural perturbation, which is relaxed in BL. The H-D unexchangeable X-H stretch weakens the hydrogen bond in Batho, but strengthens it in BL. Protein-bound water molecules that form an extended water cluster near the retinal chromophore change hydrogen bonds differently for Batho and BL, the latter containing stronger bonds than in the initial state. In addition to their structural dynamics, the present FTIR spectra at 163 K show no signals of protonated carboxylic acids, suggesting that E181 is deprotonated.

## ACKNOWLEDGEMENTS

This work was financially supported by grants from the Japanese Ministry of Education, Culture, Sports, Science and Technology to H.K. (25104009, 15H02391).

## Author Contributions

H. K. directed the research, and wrote the manuscript. S. H. prepared samples and performed all experiments with the help of K. K. S. H. and K. K. analyzed the data with H. I. All authors discussed and commented on the manuscript.

